# Genetic epidemiology of blood type, disease and trait variants, and genome-wide genetic diversity in over 11,000 domestic cats

**DOI:** 10.1101/2021.09.06.459083

**Authors:** Heidi Anderson, Stephen Davison, Katherine M. Lytle, Leena Honkanen, Jamie Freyer, Julia Mathlin, Kaisa Kyöstilä, Laura Inman, Annette Louviere, Rebecca Chodroff Foran, Oliver P. Forman, Hannes Lohi, Jonas Donner

## Abstract

In the largest DNA-based study of domestic cat to date, 11,036 individuals (10,419 pedigreed cats from 91 breeds and breed types and 617 non-pedigreed cats) were genotyped via commercial panel testing, elucidating the distribution and frequency of known genetic variants associated with blood type, disease and physical traits across cat breeds. Blood group determining variants, which are relevant clinically and in cat breeding, were genotyped to assess the across breed distribution of blood types A, B and AB. Extensive panel testing identified 13 disease-associated variants in 48 breeds or breed types for which the variant had not previously been observed, strengthening the argument for panel testing across populations. The study also indicates that multiple breed clubs have effectively used DNA testing to reduce disease-associated genetic variants within certain pedigreed cat populations. Appearance-associated genetic variation in all cats is also discussed. Additionally, we combined genotypic data with phenotype information and clinical documentation, actively conducted owner and veterinarian interviews, and recruited cats for clinical examination to investigate the causality of a number of tested variants across different breed backgrounds. Lastly, genome-wide informative SNP heterozygosity levels were calculated to obtain a comparable measure of the genetic diversity in different cat breeds.

This study represents the first comprehensive exploration of informative Mendelian variants in felines by screening over 10,000 domestic cats. The results qualitatively contribute to the understanding of feline variant heritage and genetic diversity and demonstrate the clinical utility and importance of such information in supporting breeding programs and the research community. The work also highlights the crucial commitment of pedigreed cat breeders and registries in supporting the establishment of large genomic databases that when combined with phenotype information can advance scientific understanding and provide insights that can be applied to improve the health and welfare of cats.

## INTRODUCTION

Precision or genomic medicine, encompassing understanding of individual genetic variability in disease risk and the customization of healthcare by patient subgroups, is now feasible and likely to become a future standard-of-care in veterinary medicine and care of companion animals, including cats (1). Fueling this development for felines specifically are improved genomic resources and technologies, combined with a steady increase in the discovery of Mendelian disease and trait variants (2,3). Genetic testing is a form of precision/genomic medicine when used for diagnosing diseases or traits of clinical relevance, as the results are patient-specific and can potentially be used to tailor treatment to the disease and the patient (4). Direct-to-consumer genetic testing is now readily available, and further empowers owners to proactively invest in precision medicine for their pet.

There is also an opportunity for the insight gained from feline population-based genomic data to have a transformational impact on felines and humans (1). Domestic cats are valued companions to humans, with owners seeking high quality veterinary care to support long-term wellness and positive outcomes for disease treatment. Cat genomes are structured in a similar way to humans, with 90% of their genes having a human homologue; and cats and humans also suffer from many similar diseases (1). Domestic cats, like dogs, can serve as naturally occurring models for many human diseases, in which the development of therapeutic treatments may be helpful for both veterinary and human patients (1,5–9).

While domestication of cats approximately 10,000 years ago did not introduce drastic genomic changes compared with undomesticated Felidae (10), pedigreed cats emerged about 150 years ago and do represent small genetically unique subpopulations of domestic cats. The genetic research of isolated populations in humans, dogs, and more recently cats has vastly improved identification of genetic disease and trait variants for use in genetic diagnostics and precision medicine. As an example, researchers have identified specific genetic variants of the *CMAH* gene that determine blood type in different cat breeds (11–15). Research on genetically isolated populations has further highlighted that one of the consequences of isolation -reduced genetic diversity- can manifest as an increase in the number of health conditions that are identified (16–19). For more than a decade, it has been common practice to eradicate disease-associated variants from pedigreed cat breeding populations using DNA testing. However, the focus on eradicating single DNA variants from a breed could contribute to severe loss of genetic diversity, especially if implemented strictly instead of thoughtfully (20).

Our previous work has elucidated that comprehensive screening of genetic variants in dogs is convenient and justified as it provides information to support breeding programs, veterinary care and health research (21). Further large-scale multiplex screening approaches were taken to characterize canine disease heritage and the relative frequency and distribution of disease-associated variants across breeds (22), and to explore the frequency of known canine appearance-associated variants among dog breeds (23). As we will demonstrate, such efforts are now equally feasible in cats and hold comparable promise for gaining insight into the genetic epidemiology of feline diseases and traits to better inform feline breeding decisions and establish the foundation for precision medicine of individuals, populations, and breeds.

This study represents the first comprehensive genetic evaluation of known feline disease and trait variants through the examination of 80 variants in 10,419 pedigreed cats and 617 non-pedigreed cats. These results provide a first glance into feline mutation heritage across nearly 100 breeds and breed types, and underscore the importance of large-scale population screening studies in improving veterinary diagnostics, breeding programs, and health recommendations for all cats.

## Materials and Methods

### Study Sample Population

The cat study population consisted of 10,419 pedigreed cat samples and 617 non-pedigreed cats, which was obtained during the development and provision of the commercially available MyCatDNA™ and Optimal Selection™ Feline tests (Wisdom Panel, Helsinki, Finland and Wisdom Panel, Vancouver, WA, USA, respectively) between 2016 and 2021. The 10,419 pedigreed cat samples represented 91 breeds and breed varieties, with 84 (89.3%) breeds and varieties represented by 5 or more individuals (S1 Table).

The breed of a cat was reported by its owner with accompanying registration information confirming the cat was registered with The International Cat Association (TICA) Fédération Internationale Féline (FiFe), The Cat Fanciers’ Association (CFA), or World Cat Federation (WCF) standards. A few additional breeds not yet recognized by any major breed registry but with an established community of breed hobbyists were also considered breeds for the purposes of this study. The non-pedigreed cat sample set consisted of mixed breeds; breed crosses or random-bred cats. The tested cats were most often from the United States of America (54.9%), while cats from Finland (17.4%), Canada (5.3%), United Kingdom (3.5%), Norway (3.5%) and Sweden (3.3%), Russia (2.5%) and France (1%) represented other notable subgroups (>1% of the sample).

### Ethics Statement

Feline DNA was obtained by Wisdom Panel as owner submitted, non-invasive cheek swab samples, and as cheek swab and blood samples collected at certified veterinary clinics in accordance with international standards for animal care and research. All cat owners provided consent for the use of their cat’s DNA sample in scientific research. University’s biobank samples were collected under the permit ESAVI/6054/04.10.03/2012 by the Animal Ethics Committee of the State Provincial Office of Southern Finland, Hämeenlinna, Finland.

### Microarray development

A custom genotyping microarray for selected feline disease and trait associated variants (S2 Table) was developed based on the robust and widely utilized Illumina Infinium XT platform (Illumina, Inc., San Diego, CA, USA), commercially available as the Complete for Cats™ / MyCatDNA™ / Optimal Selection™ Feline tests. The microarray was designed and validated for use following the same protocol and principles as previously described for canines (21). Firstly, public databases (3) and searches of the scientific literature were used to identify likely causal variants for feline Mendelian disorders and traits. Secondly, genotyping assays for the identified variants were designed according to the manufacturer’s guidelines (Illumina, Inc.). At least three technical replicates of each target sequence were included in the array design. Thirdly, the validation of each specific disease and trait assay was completed with the use of control samples of known genotype, or synthetic oligonucleotides in the case of rare conditions for which no control samples were available. In addition, owner-provided photographs contributed to phenotypic validation of trait variant tests.

### Genotyping

Microarray genotyping analyses were carried out following manufacturer-recommended standard protocols for the Illumina XT platform (Illumina, Inc.). All genotype data from samples with call rates below 98% of the analyzed markers were discarded in order to ensure high quality data, and all disease-associated variant calls were confirmed by manual review. Selected disease-associated variant findings were genotyped using standard capillary sequencing on an ABI3730xl DNA Analyzer platform (Thermo Fisher Scientific, Waltham, MA, USA) as a secondary technology to provide further validation of results that were unexpected for the breed. Sequencing was performed at the Sanger Sequencing Unit of the Finnish Institute of Molecular Medicine (FIMM). The DNA extractions and PCR-product preparation and purification were carried out as previously described in detail (21) using ~20 ng of genomic template DNA and an Amplitaq Gold Master Mix-based protocol according to the manufacturer’s instructions (Applied Biosystems, Waltham, MA, USA). The heterozygosity (*Hz*) values for each individual and breed were calculated from the genotypes of 7,815 informative SNPs that differentiate between individuals and evenly spaced informative SNPs with a median intermarker distance of ~312 kilobases.

### Clinical validations

Medical history information for genetically affected cats was collected through interviews with cat owners and veterinary clinicians. Clinical examinations were performed for confirmation of a Factor XII deficiency diagnosis by collecting a blood sample for an activated partial thromboplastin time screening test through IDEXX Laboratories (IDEXX Europe B.V., Hoofddorp, The Netherlands). Progressive Retinal Atrophy diagnosis was confirmed via ophthalmic examination performed by an ECVO (European College of Veterinary Ophthalmologists) board-certified veterinary specialist. A set of blood type determination data was available through the records of the former Genlab (Niini, Helsinki, Finland). All additional phenotype information (clinical or trait) and documentation was obtained through voluntary owner submissions.

## Results

### Overview of genotyping results

A total of 11,036 domestic cats, mainly pedigreed cats (N=10,419) representing 91 breeds and breed types and an additional 617 non-pedigreed cats, were genotyped using a custom Illumina XT microarray. Genetic screening included genotyping of 7,815 informative SNP markers across the genome and 82 variants associated with blood type, diseases, and/or physical appearance; 78 of which were evaluated in the entire 11,036 cat study sample and four more recently included variants were screened for in a subset of 2,186 samples (19.8% of the study sample; S1 Table, and S2-S3 Tables). We observed 51 (62.1%) of the 82 tested variants at least once in this study cohort, including 22 (44.9%) disease-associated variants and 29 (56.9%) variants associated with an appearance-associated trait or blood type (Table 1, S3 Table). The maximum number of disease-associated variants observed in any one individual was four. We observed that 2,480 (22.5%) of the tested cats had at least one disease-associated variant present, and 452 (4.1%) of the tested cats were potentially at risk for at least one health condition, in accordance with the disorder modes of inheritance. Of the disease-associated variants, 19 were observed in pedigreed cats, and 12 were also found in non-pedigreed cats. Three disease-associated variants were solely observed in non-pedigreed cat samples (Fig 1).

**Table 1.** Tested disease-associated variants and their frequencies in all cats.

| Tested Disease-associated Variant | MOI | Gene | Variant Details | Rank | Variant Allele Frequency (%) |
| --- | --- | --- | --- | --- | --- |
| Factor XII Deficiency | AR | *F12* | c.1631G>C | 1. | 7.01%* |
| Pyruvate Kinase Deficiency (PK-def) | AR | *PKLR* | c.693+304G>A | 2. | 2.94 % |
| Factor XII Deficiency | AR | *F12* | c.1321delC | 1. | 1.34 % |
| Progressive Retinal Atrophy (rdAc-PRA) | AR | *CEP290* | IVS50 + 9T>G | 4. | 1.13 % |
| Progressive Retinal Atrophy | AR | *KIF3B* | c.1000G>A | 5. | 1.11 % |
| Hypertrophic Cardiomyopathy; HCM (A31P) | AD | *MYBPC3* | c.91G>C | 6. | 0.85 % |
| MDR1 Medication Sensitivity | AR | *ABCB1* | 11930_1931del TC | 7. | 0.58 % |
| Osteochondrodysplasia and Earfold | AD | *TRPV4* | c.1024G>T | 8. | 0.42 % |
| Hypertrophic Cardiomyopathy; HCM (R820W) | AD | *MYBPC3* | c.2460C>T | 9. | 0.20 % |
| Spinal Muscular Atrophy | AR | *LIX1 -LNPEP* | a ~140 kb partial gene deletion | 10. | 0.06 % |
| Polycystic Kidney Disease (PKD) | AD | *PKD1* | c.10063C>A | 11. | 0.05 % |
| Congenital Myasthenic Syndrome | AR | *COLQ* | c.1190G>A | 12. | 0.05 % |
| Cystinuria Type B | AR | *SLC7A9* | c.881T>A | 13. | 0.05 % |
| Hyperoxaluria Type II | AR | *GRHPR* | point mutation, G to A, at the 3# splice acceptor site of intron 4 | 14. | 0.04 % |
| Familial Episodic Hypokalemic<br>Polymyopathy | AR | *WNK4* | c.2899C>T | 15. | 0.02 % |
| Lipoprotein Lipase Deficiency | AR | *LPL* | c.1234G>A | 16. | 0.02 % |
| Burmese Head Defect | AR | *ALX1* | c.496delCTCTCAGGACTG | 16. | 0.02 % |
| GM2 Gangliosidosis Type II | AR | *HEXB* | c.1244-8_1250del15 | 18. | 0.01 % |
| Glycogen Storage Disease | AR | *GBE1* | g. IVS11+1552_IVS12-<br>1339 del6 | 18. | 0.01 % |
| Myotonia Congenita | AR | *CLCN1* | c.1930+1G>T | 20. | 0.005 % |
| Acute Intermittent Porphyria (5<br>variants) | AD | *HMBS* | c.842_844delGAG,<br>c.189dupT, c.250G>A,<br>c.107_110delACAG,<br>c.826-1G>A |  | NA |
| Autoimmune Lymphoproliferative<br>Syndrome | AR | *FASL* | c.413_414insA |  | NA |
| Congenital Adrenal Hyperplasia | AR | *CYP11B1* | Exon 7 G>A |  | NA |
| Congenital Erythropoietic Porphyria | AR | *UROS* | c.331G>A + c.140C>T |  | NA |
| Cystinuria Type 1A | AR | *SCL3A1* | c.1342C>T |  | NA |
| Cystinuria Type B (2 variants) | AR | *SCL7A9* | c.706G>A, c.1175C>T |  | NA |
| Dihydropyrimidinase Deficiency | AR | *DPYS* | c.1303G>A |  | NA |
| Glutaric Aciduria Type II | AR | *ETFDH* | c.692T>G |  | NA |
| GM1 Gangliosidosis | AR | *GLB1* | c.1448G>C |  | NA |
| GM2 Gangliosidosis | AR | *GM2A* | c.390_393GGTC |  | NA |
| GM2 Gangliosidosis Type II (2 variants) | AR | *HEXB* | c.1467_1491inv,<br>c.667C>T |  | NA |
| Hemophilia B (2 variants) | X-linked | *F9* | c.247G>A, c.1014C>T |  | NA |
| Hypotrichosis | AR | *FOXN1* | c.1030_1033delCTGT |  | NA |
| Mucopolysaccharidosis Type I | AR | *IDUA* | c.1107_1109delCGA /<br>c.1108_1110GAC |  | NA |
| Mucopolysaccharidosis Type VI | AR | *ARSB* | c.1558G>A |  | NA |
| Mucopolysaccharidosis Type VII (2<br>variants) | AR | *GUSB* | c.1052A>G, c.1421T>G +<br>c.1424C>T |  | NA |
| Progressive Retinal Atrophy | AR | *AIPL1* | c.577C>T |  | NA |
| Sphingomyelinosis; Niemann-Pick C1 | AR | *NPC1* | c.2864G>C |  | NA |
| Sphingomyelinosis; Niemann-Pick C2 | AR | *NPC2* | c.82+5G>A |  | NA |
| Vitamin D-Dependent Rickets | AR | *CYP27B1* | c.637G>T |  | NA |
| **Potential Non-causal Variant<sup>#</sup>** |  |  |  |  |  |
| Mucopolysaccharidosis Type VI Modifier |  | *ARSB* | c.1427T>C |  | 2.51 % |
| Congenital Erythropoietic Porphyria |  | *UROS* | c.140C>T |  | 0.74 % |

\*The frequency value is based on a subset of 2,186 samples (19.8% of the full study sample) screened for this variant. # Genetic screening for these variants is available, but the predictive value of disease is low. Abbreviations: MOI; mode of inheritance, AR; autosomal recessive, AD; autosomal dominant, NA; not applicable.

**Fig 1.**
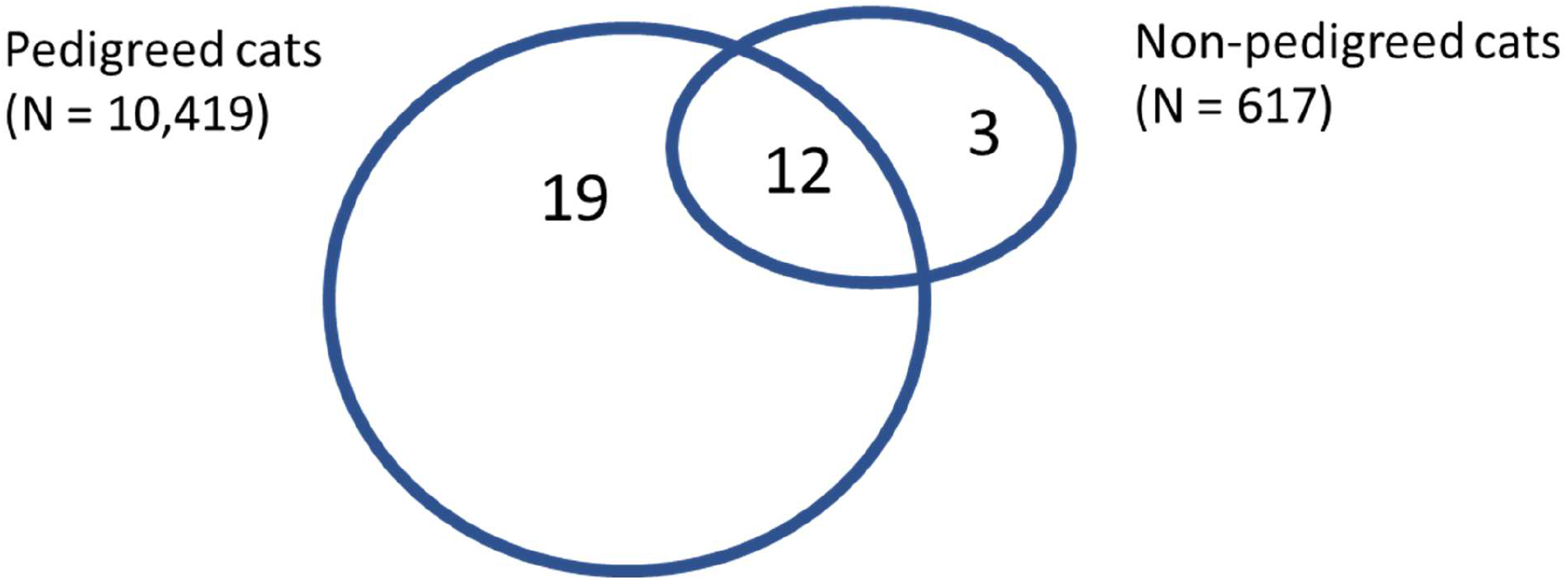
Distribution of disease-associated variants within pedigreed and non-pedigreed cat populations.

While we observed several disease-associated variants in the breeds with documented occurrence, we also detected 17 disease-associated variants in additional breeds or breed types with no previously reported frequency. These disease-associated variant discoveries and frequencies in additional breeds are listed in Table 2.

**Table 2.**
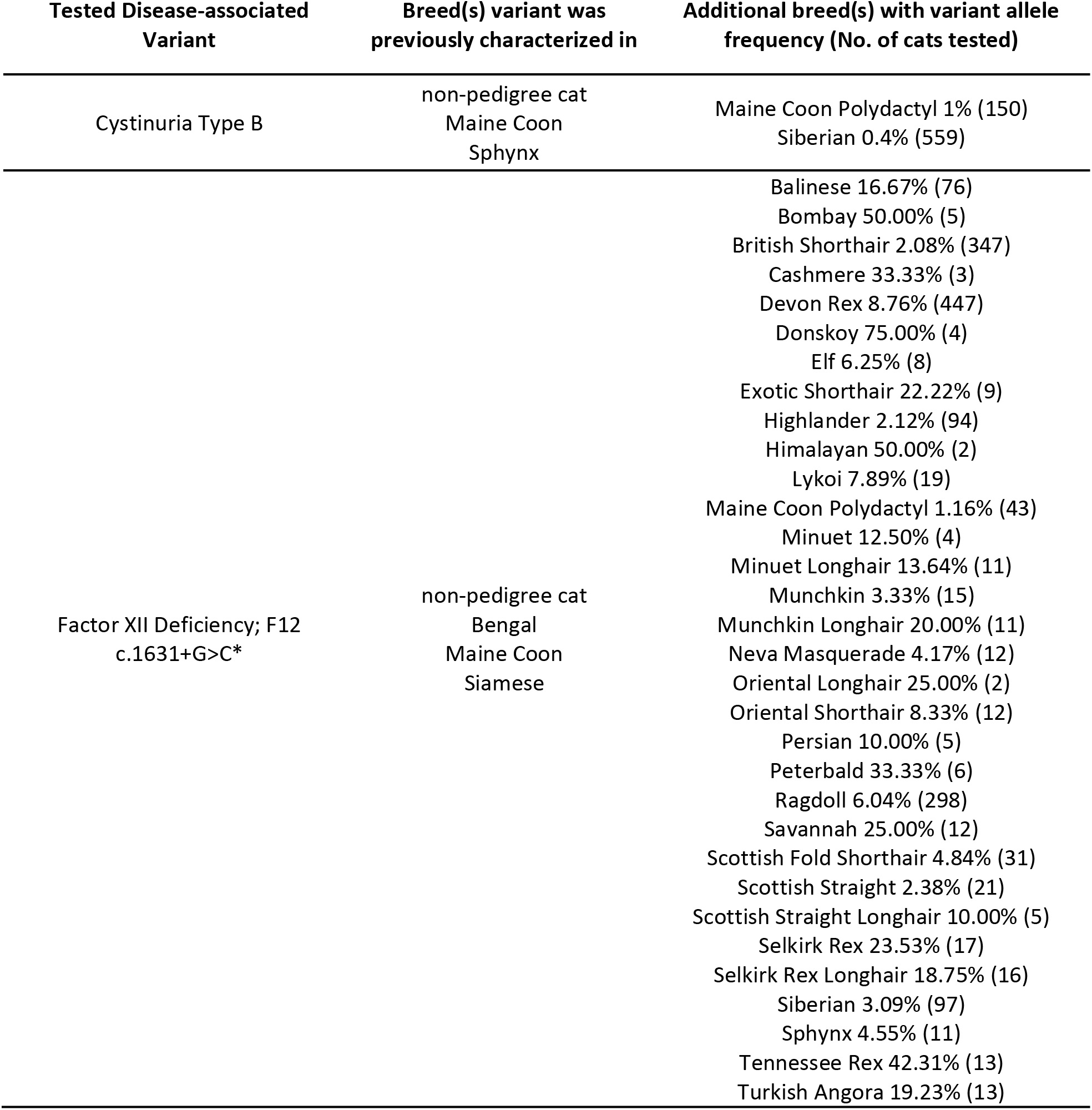
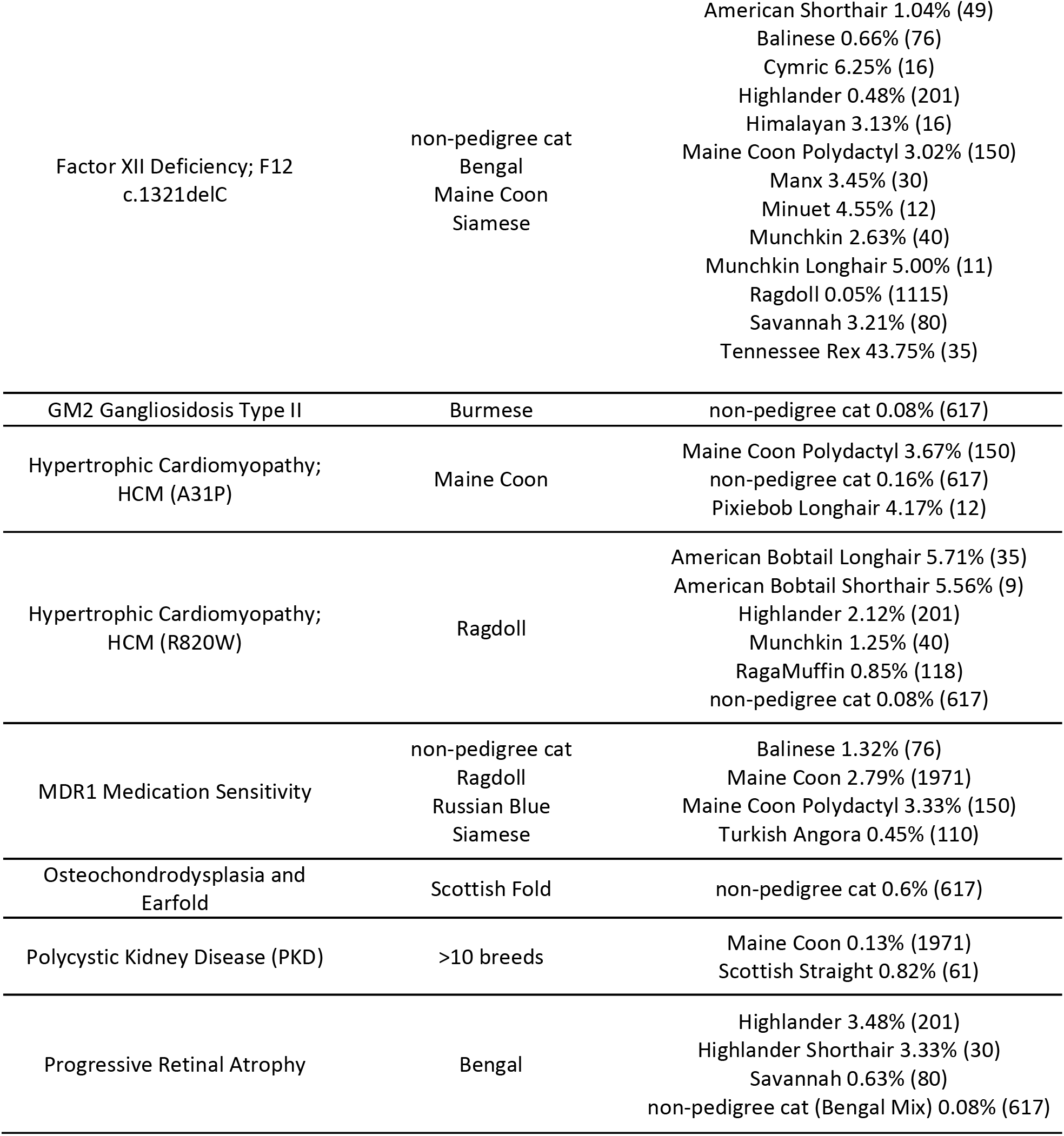
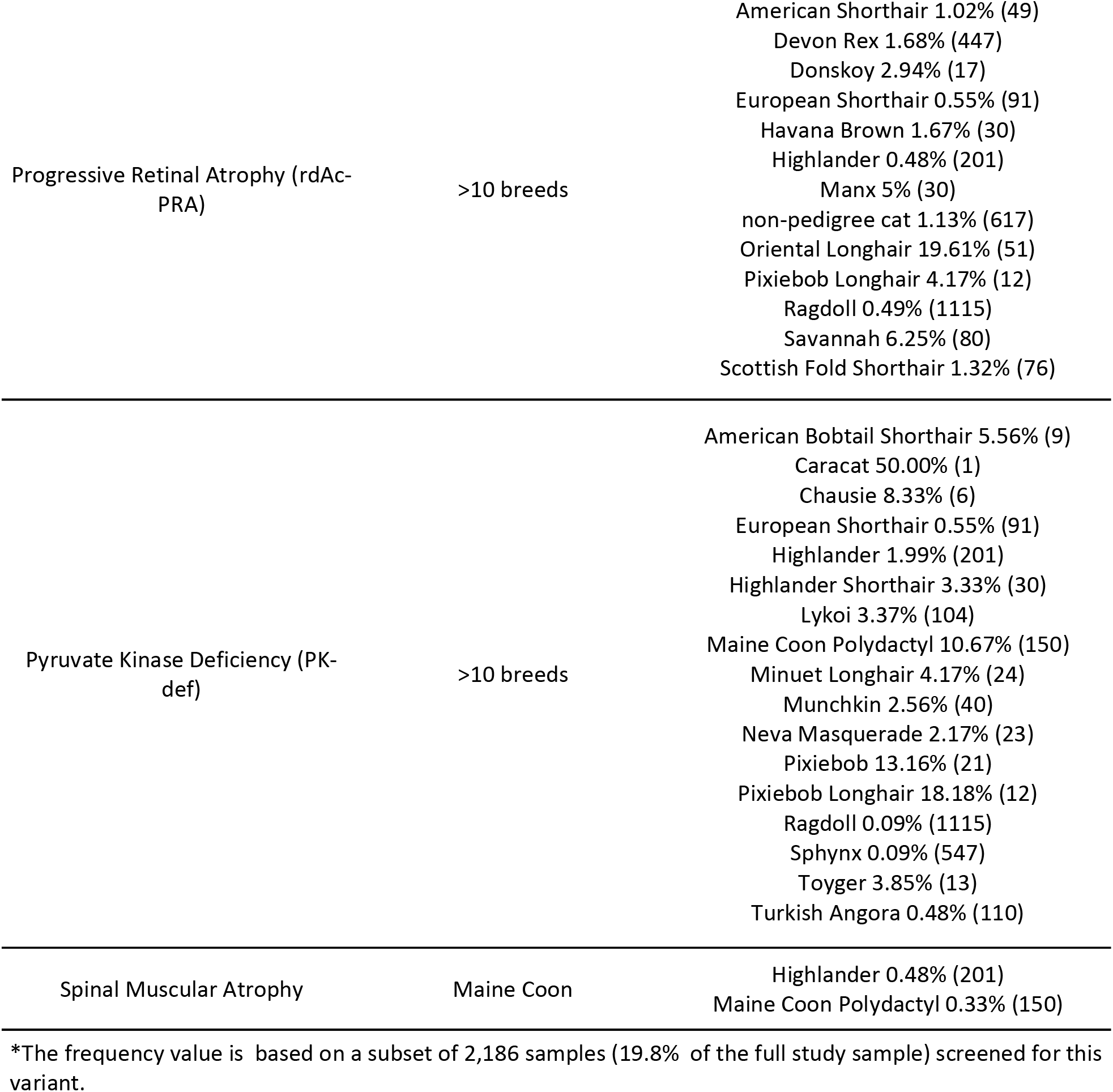
Summary of disease-associated variant findings in additional breeds.

In this study, we also investigated the tested variants’ significance in additional breeds by combining genotype data with information obtained from cat owners and/or veterinary clinicians through a variety of means, including voluntarily submitted photographs, submission of clinical documentation, or interviews with a Wisdom Panel veterinarian or geneticist. In addition, genetically affected cats were recruited for clinicopathological investigations where possible. Finally, the variant frequencies were examined in the light of genome-wide genetic diversity measured as a percentage of heterozygous SNPs.

### Genetic epidemiology of the common AB blood group system across breeds and breed types

The major feline AB blood groups including blood type A, blood type B and the rare blood type AB are caused by functional differences in the cytidine monophospho-N-acetylneuraminic acid hydroxylase enzyme encoded by the *CMAH* gene (11–15). According to current convention (2019 typing panel) (14,15), genetic testing for blood types A, B and AB should be based on panel testing of four likely causal variants of *CMAH* to assist veterinary clinicians and breeders in recognizing, confirming, and avoiding blood incompatibilities. In this study, a subset of 2,179 cats (19.7%) was genotyped for all four *CMAH* variants of the proposed genotyping scheme: common b variant 1, c.268T>A; b variant 2, c.179G>T (discovered in Turkish breeds); the c variant, c.364C>T resulting in blood type AB; and the b variant 3, c.1322delT (discovered in the Ragdoll), which was more recently added to the WISDOM PANEL genotyping platform. All 11,036 cats were genotyped for the first three variants.

We observed b variant 1, b variant 2; and the c variant to be widely distributed across breeds with frequencies of 12.6%, 1.6% and 1.5% in all cats, respectively (S4 Table). No more than two variants in total were found in any individual cat. Across the 2,179 genotyped cats representing 69 breeds and varieties, the b variant 3 was exclusively found in Ragdolls (16.9% allele frequency) and in a Ragdoll mix (S4 Table). Since the b variant 3 was found only in cats with Ragdoll ancestry, applying the assumption that the entire dataset was consistent with this finding allowed genotypic interpretations for blood type determination to be based on three variants (the b variant 1 and 2 and the c variant) in all breeds except for Ragdoll, in which a four variant-based interpretation of genotype (including the b variants 1, 2, and 3, and the c variant) was used.

Based on genetic blood type determination, blood type B was most common in the following five breeds or breed types: American Curl (40.4%), British Shorthair breed types (20.3%), Cornish Rex (33%), Devon Rex (30.3%) and Havana Brown (20%). In addition, the breeds in which the rare blood type AB was present with a frequency of >1% were European Shorthair (2.2%), Lykoi (1%), Scottish Fold (3.3%), RagaMuffin (3.4%), Ragdoll (2.2%) and Russian Blue breed types (1.5%). The proportions of type A, type B and type AB blood for each breed or breed group with >15 individuals tested are shown in Table 3 (breed groups listed in S5 Table). In this study, blood typing results from serological tests were available for 220 cats. We found that blood types assigned according to the cat’s genotype and serological tests were 99.5% concordant. One Ragdoll with c/c genotype shown to result in blood type AB (11,14,15), had been determined serologically to have blood type A according to the cat’s owner, although retesting to confirm this result or further clinical investigation was not performed.

**Table 3.**
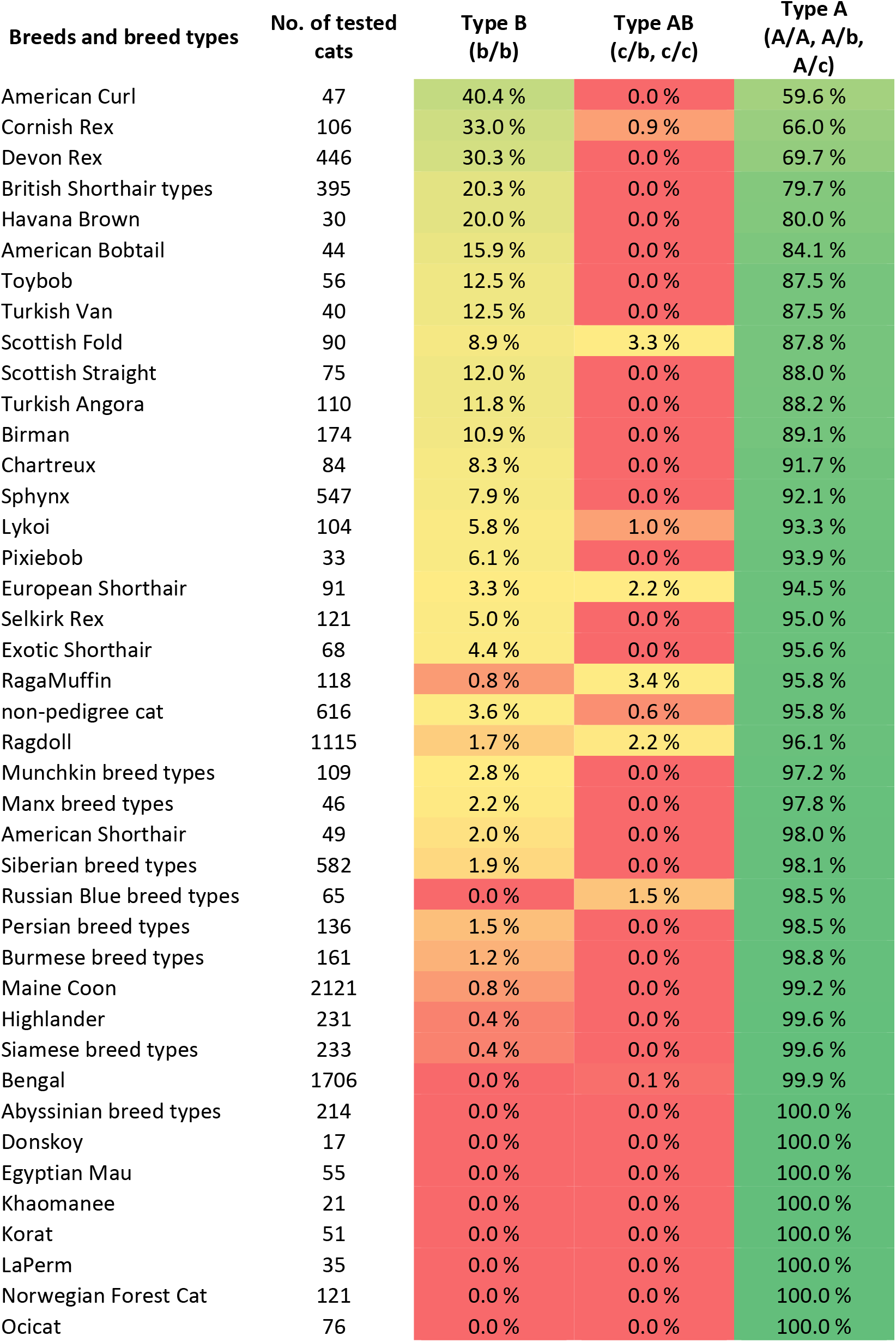
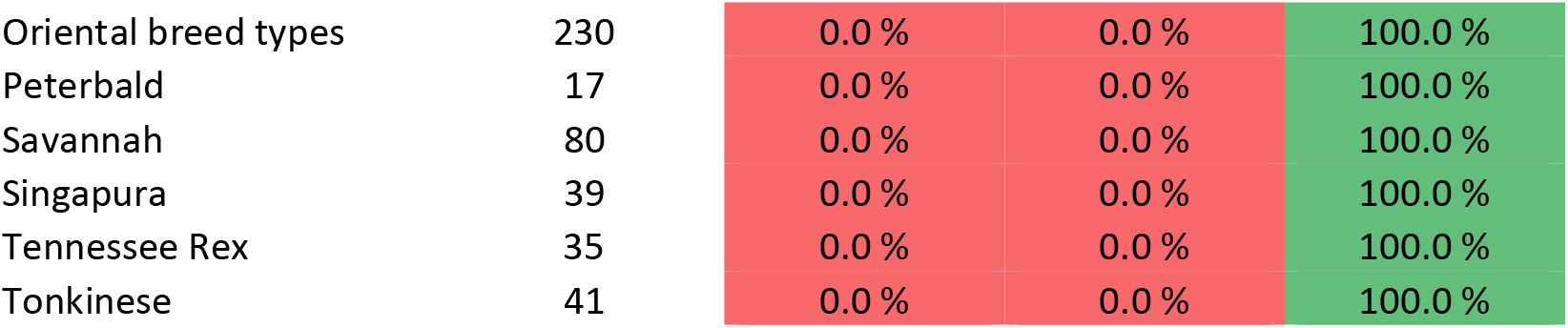
Genetically determined proportions of type A, type B and type AB blood for each breed or breed type with >15 individuals tested.

### Factor XII Deficiency and Pyruvate Kinase Deficiency are widespread blood disorders in the cat population

Factor XII Deficiency is a widely distributed heritable disorder in the domestic cat population (24). Factor XII Deficiency is a clinical hemostatic defect that manifests as a prolonged activated partial thromboplastin time (aPTT) which would be observed in a presurgical coagulation assay, but does not require transfusions (25). Two variants of the *F12* gene (c.1321delC and c.1631G>C) have been identified in a colony of inbred cats from the United States and in a litter of cats from Japan, respectively. Both variants segregate consistently with cases of Factor XII Deficiency. The variants also co-segregate, although variant c.1631G>C is likely to have originated first, as it is observed without the c.1321delC variant. These variants are both considered common in the domestic cat (25–27). The most severe aPTT prolongation is observed in cats homozygous for both variants (25). In accordance with the previous observations (25), we noted that the c.1321delC variant always co-segregates with c.1631G>C, while one or two copies of the latter can be inherited in the absence of the c.1321delC variant. The observed variant frequencies for the c.1321delC and c.1631G>C variants were 1.3% and 7% in all cats (S4 Table). The frequency of the c.1631C variant was based on a subset of 2186 genotyped cats, as the variant represented a more recent discovery added to the WISDOM PANEL genotyping platform. Notably, the c.1631G>C variant present alone without the second variant is observed at high frequency in the Donskoy (75%), Bombay (50%) and Himalayan (50%) breeds, while Tennessee Rex represents a breed with high frequency for both variants (>40%) observed together (S4 Table). In all, we discovered the two variants of *F12* gene present in 40 additional breeds. Our data confirm the presence of the identified *F12* variants in the following breeds in which cases of Factor XII Deficiency have been documented: Himalayan, Maine Coon, Manx, Munchkin, Oriental Shorthair, Persian, Ragdoll, Siberian and Siamese. The c.1321delC variant was not discovered in any of the 121 genotyped Norwegian Forest Cats, nor was the c.1631G>C variant (examined in 16 Norwegian Forest Cats) despite documented cases of Factor XII Deficiency in the breed (25). Consequently, it may be possible that these cats are affected by other candidate variants of *F12* such as the c.1549C>T variant, which was recently identified (25), but not screened in this study. Furthermore, we could not confirm the tested variants’ presence in the Turkish Van. We, therefore, could not corroborate the documented association with Factor XII Deficiency (25), though we did discover the presence of the c.1631G>C variant in the related Turkish Angora breed. Finally, we obtained laboratory results from a 10-month-old intact female Maine Coon homozygous for the c.1321delC and c.1631G>C variants of *F12* gene. This individual showed prolonged aPTT of >180 seconds (laboratory reference <13.4 seconds), confirming Factor XII Deficiency in the Maine Coon and the association of the tested variants with clinical signs.

Pyruvate Kinase Deficiency (PK-def) is an inherited anemia characterized by low levels of the pyruvate kinase enzyme. This insufficient presence of the PK enzyme causes red blood cells to break easily, resulting in hemolytic anemia. A single nucleotide substitution variant (c.693+304G>A) of intron 5 in the genomic DNA splice site of the *PKLR* gene has been associated with the manifestation of PK-def in the Abyssinian and Somali breeds (28), but has also been previously identified in at least 15 additional cat breeds. PK-def associated with the identified variant has marked clinical variability, including variation in the age of disease onset and disease severity (28,29). Here we report the presence of the *PKLR* variant in an additional 35 breeds and breed types (Table 1). Where the breed was represented by at least ten individuals in our study set, the breeds with the highest allele frequencies were the Pixiebob (16.6%), Pixiebob Longhair (13.5%), Egyptian Mau (12.7%) and Maine Coon Polydactyl (10.5%). The updated *PKLR* variant frequencies for the Abyssinian and Somali breeds were 3.1% and 2.2%, respectively (S4 Table). Several young cats from Bengal, Maine Coon and Maine Coon Polydactyl breeds had two copies of the PK-def associated variant and are therefore clinically at risk. In the scope of this study, we were able to interview the owner of a 3-year-9-month-old female Bengal cat. The owner reported a veterinarian had obtained a complete blood count (CBC) for this cat, however, the test results at that time did not reveal the presence of anemia. Additionally, we reached out to several breed clubs to ask whether clinical PK-def has been documented in the Maine Coon, but discussions were inconclusive. Further clinical data are required to confirm clinical signs manifesting in the additional breeds identified with the *PKLR* variant.

### Panel screening reveals autosomal dominant disease-associated variants affecting cats in additional breeds

We screened for four feline disease-associated variants that most closely follow an autosomal dominant mode of inheritance in clinical settings: Polycystic Kidney Disease (PKD) (30), two Hypertrophic Cardiomyopathy (HCM) variants (31,32), and Osteochondrodysplasia and Earfold (33).

Polycystic Kidney Disease (PKD) is a severe autosomal dominant (homozygous lethal) condition in which clusters of cysts present at birth develop in the kidney and other organs, causing chronic kidney disease which can lead to kidney failure (34,35). PKD is caused by a stop codon in exon 29 of *PKD1* (c.10063C>A), resulting in a truncated form of the gene, which was discovered in Persian cats with ~40% frequency in the Persian cat population worldwide (36–39). Genetic testing was introduced into the breeding programs of Persians and some Persian-related cats. We report that the overall frequency of the *PKD1* variant in the screened breeds has reduced notably from what was previously reported (S4 Table). The *PKD1* variant was identified in the Maine Coon, a breed in which it had not been previously documented in the peer-reviewed literature. Clinical manifestation of PKD in a genetically affected female Maine Coon, diagnosed at the age of 3 months, was confirmed after interviewing the cat’s owner and assessing associated diagnostic documentation including an ultrasound of the kidneys in which numerous, round, well-defined cysts were observed bilaterally throughout the renal cortex and medulla. This finding provides confirmation that genetic screening for the *PKD1* variant in the Maine Coon is a clinically relevant marker for PKD.

Hypertrophic cardiomyopathy (HCM) is the most common heart disease in domestic cats. Two independent condition predisposing variants of the *MYBPC3* gene c.91G>C p.(A31P) and c.1024G>T p.(R820W) have been associated with HCM in Maine Coon and Ragdoll breeds, respectively (31,32,40). In the heterozygous state, the likelihood of developing clinical HCM early in life is very low. However, supporting the autosomal dominant mode of inheritance, regional diastolic and systolic dysfunction has been observed in heterozygous asymptomatic cats (41,42). In the homozygous state, the development of HCM (A31P) is highly likely in the Maine Coon with risk increasing with age (43). Similarly, in Ragdoll cats the (R820W) heterozygous cats have a normal life expectancy, while homozygous cats are likely to have a shortened life span (44). In the present study, we found higher frequencies of HCM (A31P) and HCM (R820W) variants present in additional breeds (Table 1). The observed cats were young, and all were carrying one copy of either of the variant alleles; no further clinical validations were pursued in the scope of this study. Yet, given the association of these likely causal variants with HCM, these variants suggest potential molecular explanations for cases of HCM in these additional breeds.

Osteochondrodysplasia and Earfold is a highly penetrant autosomal dominant condition caused by a missense variant (c.1024G>T) in the *TRPV4* gene resulting in congenital degenerative osteochondrodysplasia or “Scottish Fold Syndrome”, which results in skeletal deformities such as a short, thick, inflexible tail and malformation of the distal fore- and hindlimbs, which can lead to a stilted gait (33). We observed at least one copy of the *TRPV4* variant in all 90 tested Scottish Fold cats, and the *TRPV4* variant was absent in all 75 Scottish Straight cats tested. We also discovered one copy of the *TRPV4* variant in a crossbred cat resulting from the mating of a Scottish Fold and a Highlander. As the ear phenotype of the kitten was curled-back as presented in the Highlander breed, rather than folded forward as seen in the Scottish Fold, observation of the *TRPV4* variant was not entirely expected. However, the kitten did present a stiff and inflexible shortened tail, characteristic of *TRPV4* variant carriers. It therefore would appear that the yet unknown variant that is causing the Highlander ear type masks the Scottish Fold ear phenotype caused by the *TRPV4* variant, when the two variants are inherited together. In another recent study, a cat registered as an American Curl with curled ears was diagnosed with osteochondrodysplasia and genotypically showed one copy of *TRPV4* variant (45). Thus, we report a second case of Osteochondrodysplasia in which the cat’s ear phenotype belied the presence of the causal variant.

### Molecular heterogeneity of feline hereditary retinal dystrophies

The disease-associated variants for retinal dystrophies screened in this study include *CEP290*, *KIF3B* and *AIPL1*. The *CEP290* variant associated with late-onset Progressive Retinal Atrophy (rdAc-PRA) is present in many pedigreed breeds (46,47); here we document a frequency of 1.1% in all cats. We have identified the presence of the *CEP290* variant in 20 additional breeds. Some of the highest *CEP290* variant frequencies were observed in the Peterbald (26.5%) and in one of the additionally identified breeds, the Oriental Longhair (19.6%). The variant of the *KIF3B* gene, recently associated with an early-onset Progressive Retinal Atrophy in the Bengal (48), was present in 6.9% of the Bengal breed (resulting in a frequency of 1.1% in all cats) (Table 1–2, and S4 Table). This variant was additionally discovered in the Highlander breed types and the Savannah. The *AIPL1* variant associated with Progressive Retinal Atrophy (discovered in the Persian) was the rarest variant associated with retinal dystrophies (49), that was screened in a subset of 2,186 samples and not observed at all (Table 1, S4 Table). To pursue clinical validation of our findings, we recruited a 10-year-3-month-old female Oriental Longhair, homozygous for *CEP290*. Clinical validation was initiated with an owner interview, in which the owner reported no apparent changes in the cat’s behavior that were suggestive of vision loss. Yet, during an ophthalmic examination, a marked discoloration of pigmentation of the tapetal fundus with a slight vascular attenuation was noted, confirming the presence of retinal degeneration. We demonstrate that rdAc-PRA does manifest clinically in the Oriental Longhair. Some vision may be retained longer than the previously reported 3-7 years (5), at least for this particular breed example.

### Feline MDR1 Medication Sensitivity associated with adverse medication reactions in the Maine Coon

Feline MDR1 Medication Sensitivity is a disorder associated with severe adverse reactions after exposure to medications that use the p-glycoprotein drug transporter. This genetic condition is caused by a two base pair deletion within exon 15 of the *ABCB1* gene resulting in abnormal p-glycoprotein (50). While functional p-glycoprotein plays a significant part in the blood-brain barrier that prevents various drugs and chemicals in the bloodstream from entering the brain, a defective p-glycoprotein allows more drugs to cross this barrier, thus increasing the neurological effects of some medications. Severe macrocyclic lactone-induced neurologic toxicosis has previously been reported in cats homozygous for the MDR1 variant receiving either a subcutaneously administered dose of ivermectin or a topically administered eprinomectin-containing antiparasitic product labeled for cats (50,51). We report the frequency of the *ABCB1* variant as 0.6% of all genotyped cats, in addition to the discovery of the variant in the Maine Coon, Ragdoll, Siamese and Turkish Van breeds (Tables 1–2; S4 Table). Our veterinarians interviewed three owners of cats identified as homozygous for the MDR1 variant to assess their cat’s medical history. The cats consisted of a 1-year-4-month-old intact female Maine Coon, a 2-year-3-month-old intact female Ragdoll, and a 3-year-2-month-old male Maine Coon. Only one of the cats had undergone anesthesia, and reportedly showed a delayed recovery with mild lethargy documented on the following day, but fully recovered. All three cats had been administered topical flea medications (of varying brands) with no discernable side effects, however, none of the medications applied contained eprinomectin.

### Genetic diagnosis plays a crucial role in the diagnosis of uncommon inherited disease

We identified a cat genetically affected with Myotonia Congenita, an uncommon recessively inherited disorder manifesting as an inability of the muscles to relax after contraction, which is caused by a variant in the *CLCN1* gene (52). This is a sporadic condition that was discovered in a rescue domestic cat population in Winnipeg, Canada. While the variant was not identified in any pedigreed cats (which make up a large proportion of the study sample), we discovered two copies of the variant in a single non-pedigreed domestic cat in Oregon, United States. In the owner interview, we learned that the genetic diagnosis was crucial in assisting with clinical diagnosis. While this is an incurable condition, having the correct diagnosis helps ensure that the cat is getting appropriate supportive care. The owner confirmed that this cat, initially misdiagnosed with flea bite anemia at the age of 13 weeks, has a disease manifestation that includes fainting spells when startled, prolonged prolapse of the nictitating membrane, hypertrophic musculature, flattened ears, motor dysfunction, shortened gait, and limited range of motion in the jaw. The cat also shows a characteristic “smile” after a yawn or a meow due to delayed relaxation of the muscle in the upper lip as well as commonly has the paws protracted (Fig 2A). However, the owner mentioned that this cat, currently 4-year-2-month-old, is not drooling or showing any dental defects, which differs from the original disease description (52).

**Fig 2.**
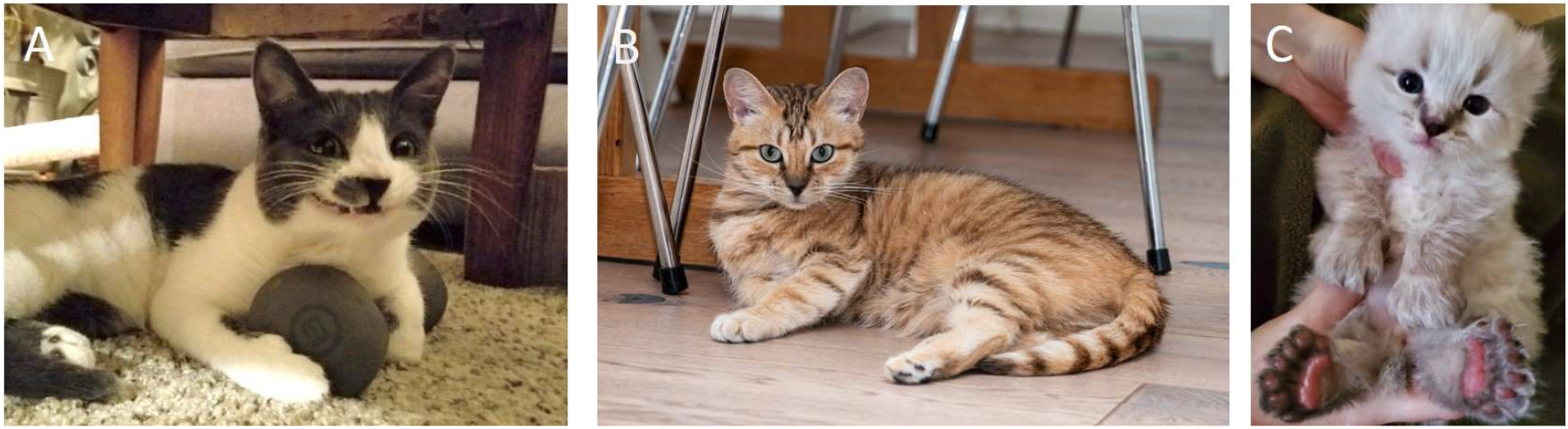
A) The signature “smile” of a cat with Myotonia Congenita; B) The rare coat color phenotype Amber in a random-bred cat from Finland; C) Polydactyly (variant 1) also associated with extra toes in all four feet.

### Panel screening enables dissociation of variants with clinical disease

In this study we determined the frequency of the c.140C>T p.(S47F) variant of the *UROS* gene to be 0.7% across the entire study sample (Table 1, S4 Table). This variant was previously discovered along with c.331G>A p.(G111S) in a cat manifesting Congenital Erythropoietic Porphyria (CEP) (53). However, we did not observe the c.331G>A variant in any of the tested cats. While the original research revealed no cats carrying solely the c.140C>T variant, functional studies have shown that c.140C>T alone does not significantly alter the protein function (53). The manifestation of CEP includes distinctively stained brownish-yellow teeth that turn fluorescent pink under UV light. Due to the high frequency of c.140C>T in some cat breeds and because some DNA testing laboratories offer tests for the two variants separately, our veterinarians reached out to the owners of cats homozygous for only the c.140C>T variant. Owners were contacted in the cases of five c.140C>T homozygous cats: a 1-year-4-month-old intact female RagaMuffin, a 1-year-5-month-old intact male RagaMuffin, a 1-year-7-month-old intact female Siberian, a 2-year-10-month-old intact female Singapura and a 3-year-7-month-old intact Toybob; none of these cats had clinical signs suggestive of CEP. Thus, the existing evidence strongly suggests that c.140C>T is a benign variant when it is not inherited along with c.331G>A. We would advise breeders and DNA laboratories not to consider c.140C>T alone a precursor for CEP.

Mucopolysaccharidosis Type VI (MPS VI), a lysosomal storage disease caused by a deficiency of *N*-acetylgalactosamine-4-sulfatase (4S), is another disease in which the roles of two variants, independently inherited in this case, have been under discussion (54). While the MPS VI variant c.1427T>C p.(L476P) of the *ARSB* gene is associated with severe disease (55), the c.1558G>A p. (D520N) variant of the *ARSB* gene is sometimes referred to as the “mild” type (56). However, it is necessary for the (D520N) variant to be inherited as a compound heterozygote with one copy of the L476P variant for the disease to manifest in its mild form. We have elected to call the D520N variant MPS VI Modifier. The D520N variant is present in large numbers of cats, with a frequency of 2.5% (Table 1, S4 Table). It is also the major allele in the Havana Brown, with an allele frequency of 76.7%. Additionally, we confirm that MPS VI (L476P) is a scarce variant, and this study cohort did not identify any cats with MPS VI variant in concordance with previously described observations (54). The high variant frequency of the benign MPS VI Modifier and very low frequency of the MPS VI variant (absent in many breeds) further confirm that breeding to avoid MPS VI should concentrate solely on managing the MPS VI variant.

### Appearance-associated variants distributed across breeds and breed types

In this study sample, the ancestral form of the appearance-associated (trait) variant was the major allele, except for the *ASIP* gene in which the derived allele a (non-agouti/solid color) showed a 56.2% variant frequency in all cats (S4 Table). The rarest derived allele was the e allele of the *MC1R* gene associated with the coat color Amber (discovered in the Norwegian Forest Cat), only observed in one random-bred cat from Finland. The observed cat was homozygous for the derived allele with pictures confirming the expression of the Amber coat color (Fig 2B).

Expectedly, the screened trait variants were present in a high number of breeds and breed types (S4 Table). Genotyping results showed overall concordance with observed phenotypes, strongly supporting the variants’ causality, with one exception where our findings are supportive of the potential additional candidate locus may play a role in the regulation of gloves phenotype (59). In Birman cats, the two adjacent missense variants of the *KIT* gene associated with breed-defining white gloving phenotype (58) were observed with a high variant frequency of 95.6% in the breed. These variants were also observed as the minor allele in a large number of breeds and breed types, and in homozygote state in some individuals of Chartreux, Highlander, Maine Coon, Ragdoll and Siberian breeds, in which they were not associated with gloving based on photographic evidence and owner reports. Of the phenotyped individuals some white areas of hair were seen in the belly or toes in (3/17) Maine Coons and (1/2) Siberian cats.

The polydactyly variant *Hw* was the most typical *LIMBR1* gene variant observed in polydactylous cats, including the Maine Coon, Pixiebob and Highlander breed types, and some non-pedigreed cats from North America. The *Hw* variant (variant 1) of *LIMBR1* was observed to be incompletely penetrant, with cats presenting with four to seven toes per paw. Higher penetrance without a more extreme phenotypic manifestation was seen in homozygous cats, which is in line with previous observations (59). However, extra toes did not manifest solely in the front feet as previously reported; photographic evidence and owner-provided details revealed the presence of extra toes on all four feet, which was formerly considered to be characteristic of the *UK1* (variant 2) and *UK2* (variant 3) variants only (60) (Fig 2C). The only cat with two copies of the *UK1* variant had two extra digits on each paw, per the owner’s description. Various owner-reported cases of polydactyl cats testing negative for the screened variants were also noted. Such cats are likely to be carrying additional variants of the *LIMBR1* gene not yet identified.

Additional variants are also likely to be associated with shortened tail phenotype. The c.5T>C variant in the gene *HES7* (discovered in the Japanese Bobtail) associated with a shortened and kinked tail (61) was observed in 100% of the Japanese Bobtail cats studied, and was also highly prevalent in the Kurilian Bobtail (96.4%) and Mekong Bobtail (66.7%), with some minor allele frequencies observed in additional bobtail breeds (S4 Table). All studied cats with one or more copies of *HES7* variant and available phenotypic information presented with a shortened tail.

The three tested variants (c.998delT, c.1169delC and c.1199delC) of the *T-box* gene (discovered in the Manx) were found in the American Bobtail, Cymric, Highland Lynx, Highlander, Manx, Pixiebob and Toybob. However, photographic evidence in these breeds revealed several short-tailed cats that were not carrying any known bobtail variant in addition to the cats that were the long-tailed versions of these breeds. The absence of known variants was especially profound in Highlander breed types and Toybob, in which cats with no known bobtail variant had a prevalence of 50-60% and 78.6%, respectively (S4 Table).

We also discovered that the c.816+1G>A variant in the *KRT71* gene which is known to result in the hairless phenotype of the Sphynx cat had a variant frequency of 74.7% in the Sphynx breed (S4 Table). Compound heterozygotes for the Sphynx c.816+1G>A variant with the Devon Rex associated curly coat variant are also hairless. A subset of 2,186 samples were genotyped for the Devon Rex curly coat in this study, however, we also identified 22 hairless Sphynx cats without any copies of the Sphynx variant suggesting that an additional unknown variant in the same or another gene entirely causes the hairless phenotype in some cats of this breed. We also confirmed that the Sphynx variant was not present in the Donskoy and Peterbald breeds, which also represent hairless phenotypes. It was recently suggested that a novel 4 base pair variant of the *LPAR6* gene identified in a Peterbald cat potentially results in a hairless coat phenotype as a homozygote or as a compound heterozygote with the c.250_253_delTTTG variant in the *LPAR6* gene which causes the curly coat of the Cornish Rex (57). We show a variant frequency of 25% for the Cornish Rex coat variant in the Donskoy, but did not identify any Cornish Rex coat variant carriers among the 17 tested Peterbald cats of this study.

### Genome-wide analysis of genetic diversity demonstrates differences between and within cat breeds

The entire data set of 11,036 samples was genotyped for 7,815 informative SNP markers distributed across the genome. In the pedigreed cat population, the median heterozygosity was 34.0% and the typical range (defined as the 10th and 90th percentile) was 27.2%-38.3%, in the non-pedigreed population, the median heterozygosity was 38.8%; and the typical range was 29.8%-41.3% (Fig 3). Of the 91 breeds tested, the median heterozygosity was calculated for 56 breeds and breed types were represented by at least 15 individuals in the dataset (S6 Table). The most diverse breeds include three of the newer cat breeds; the short-legged Munchkin, produced from a sibling mating followed by regular non-pedigree cat outcrosses (2,62); the Highlander, a crossbreed of two recent experimental hybrid cat breeds the Desert Lynx and Jungle Curl; and the Lykoi breed founded by unrelated cats expressing hypotrichosis, whose unique sparse and roaned coat phenotype may be caused by any of six different variants of the *HR* gene from six independent lineages found in four different states of the United States, Canada and France (63). The heterozygosity levels of European Shorthair, Norwegian Forest Cat, Siberian and Manx, developed from the local domestic populations with likely a larger diversity in the founder population, were above average compared with the entire pedigreed cat population. The lowest median heterozygosity measures in any pedigreed cat population were observed in the Burmese, Birman, Havana Brown, Korat, Singapura and cat breeds of the Siamese group (such as Balinese, Siamese and Oriental Shorthair), in line with previous observations (20). A full breakdown of the diversity levels per breed can be found in Supplemental Table 5.

**Fig 3.**
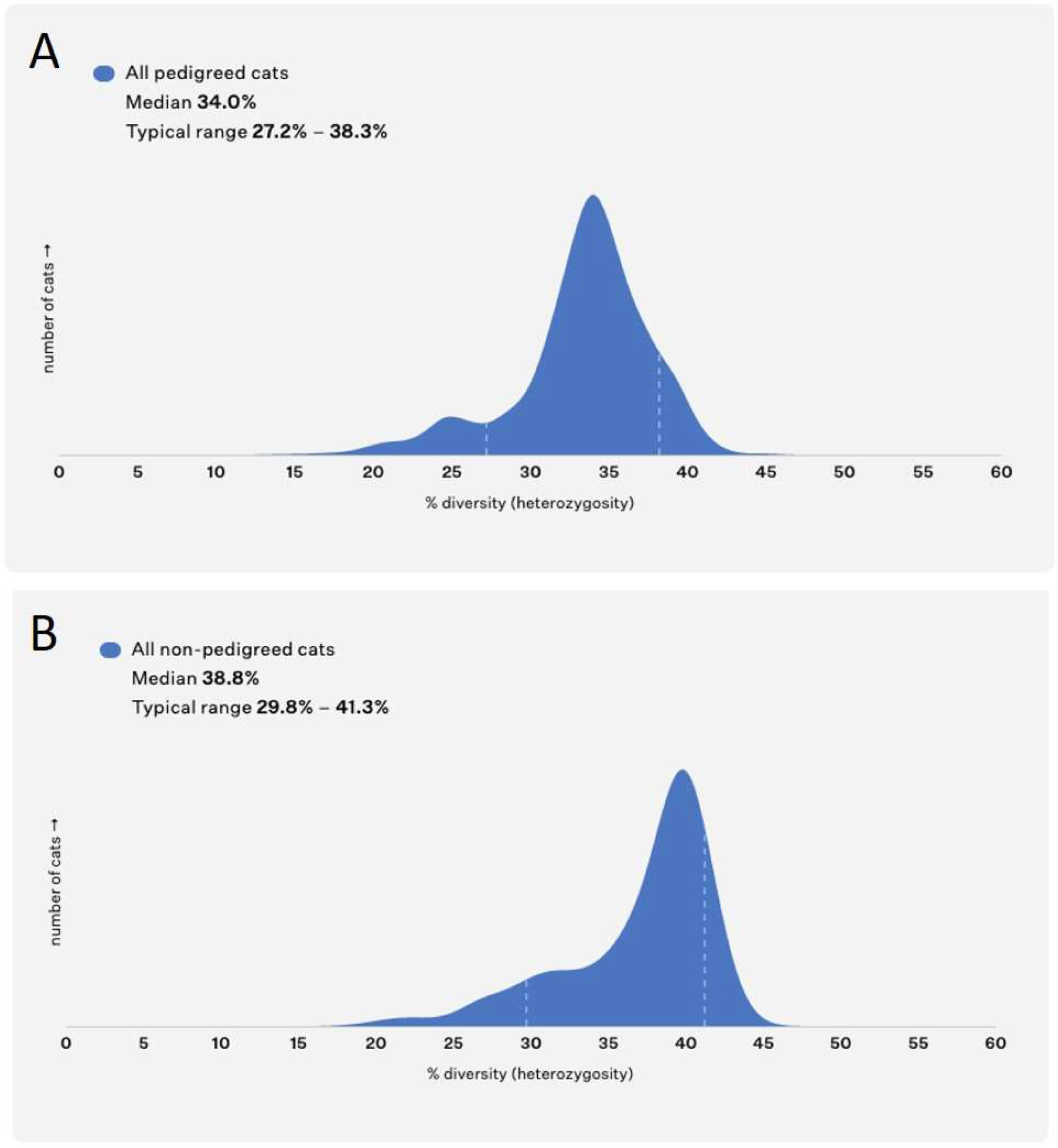
The median genetic diversity in pedigreed and non-pedigreed cat populations with typical range (the 10th and 90th percentile).

## Discussion

In the largest DNA-based feline study cohort to date, a custom genetic panel screening test was utilized to determine blood type, disease and phenotypic trait heritage, as well as the relative genome-wide genetic diversity in 11,036 domestic cats representing 91 breeds and breed types.

One specific area of focus was blood type determination, which is particularly important in cats due to its link with Neonatal Isoerythrolysis, a significant cause of fading kitten syndrome and neonatal death if the blood types of breeding pairs are not appropriately identified. We identified 17 breeds in which at least one out of ten cats in the population were blood type B. We also observed the rare blood type AB, previously described with notable prevalence only in the Ragdoll, also present in the European Shorthair, Scottish Fold breed types, and the RagaMuffin. We confirm a high concordance between blood type determination based on the proposed DNA genotyping scheme for purpose-bred cats, and serological blood type based on the 220 results available for analysis in this study. This provides further justification for the use of DNA-based blood type determination approaches in felines, provided that the appropriate carefully validated genetic variants are assayed.

Panel screening of disease-associated variants provides clinically relevant information. In addition to blood type determination, we show that the genomic data available today can assist in disease diagnosis, treatment, and preventative care. Through the comprehensive investigation of variant allele frequencies in this study cohort, we re-evaluate and provide updated variant frequency information compared to estimates provided in conjunction with the original variant discoveries. Our data suggest that disease-associated variant frequencies are now lower for many conditions (GM2 in Burmese, Hypokalemia in Burmese, HCM-A31P in Maine Coon, HCM-R820W in Ragdoll, PKD in Persian) compared to the frequencies at the time of their discovery, perhaps reflecting change over time within the breed, presumably due to genetic testing combined with informed breeding selections. We note in particular that some variants, such as rdAC-PRA, PKD, HCM-A31P and HCM-R820W appear more common in additional breeds than in the original breed in which they were discovered, likely due to lack of awareness and inadvertent selection. For example, the *PKD1* variant, initially discovered to affect nearly 40% of Persian cats, was found in this study at higher frequency in breeds with Persian background or non-related breeds than it was in the Persian breed. In fact, none of the Persian cat samples in this study were identified as carriers of the *PKD1* variant. Additional recent studies indicate that the prevalence of PKD in Persian cats of Iranian origin continues to be high (64), but also that *PKD1* is common in pedigreed and non-pedigreed cats in Japan and Turkey (65,66). Taken together, the *PKD1* variant should be seen as a potential genetic cause of PKD in any breed, such as the previously documented Neva Masquerade (67), Chartreux (54,68) or the Maine Coon as reported in this study. In this study we mainly focused on genotyping pedigreed cats and had a relatively small sample size of non-pedigreed cats. We nevertheless discovered 13 likely causal disease-associated variants in non-pedigreed cats.

This study sample provides an extensive investigation of disease-associated variant heritage across nearly 100 cat breeds. Large scale screening studies of isolated subpopulations of a species such as pedigreed cats hold great value as a secondary independent tool for validating original discoveries in as often there are made by focusing on a limited number of individuals from a single breed. Investigations that extend beyond the original discovery breed enable researchers to conclusively to understand the causal relationships between variants and diseases. For each disease-associated variant discovered in additional breeds in this study, a total of 13 different variants across tens of breeds, our follow up investigations applied a similar validation protocol as previously recommended for dogs by combining genotype information with clinical information collected and evaluated by veterinarians to assess variant manifestation in different breed backgrounds (22). These clinical phenotype evaluation studies are crucial to ensure genetic counseling information that truly offers solutions to improve the health of cats, and they are fueled by the cat community and individual breeders’ willingness to provide phenotype information and clinical documentation and participate in veterinary examinations. Here we confirm a strong relationship between several disease-associated variants (*F12*, *PKD1*, *TRPV4*, *CEP290*, *ABCB1* and *CLCN1*) and their clinical manifestations. We further suggest investigating the role of the two *MYBPC3* variants in contributing to HCM in additional breeds, and raising awareness that PK-def is a genetically common condition that may result in highly variable clinical signs and perhaps be similarly underdiagnosed in cats as it is in humans (69). Finally, after evaluation, we report two associated disease variants that have little value as markers for genetic disease. Both the c.140C>T variant of the *UROS* gene previously co-segregating with a second variant associated with CEP and the c.1558G>A variant of *ARSB* associated with mild MPS VI disease as a compound heterozygote with the c.1427T>C variant (severe disease-associated variant) were found to be the major variants in some breeds without any health impact. We advise DNA testing laboratories to discontinue offering a test for CEP based solely on the use of the c.140C>T variant. Moreover, similar to previous investigations (54), we further emphasize that prevention of MPS VI should focus entirely on managing the c.1427T>C severe variant in the cat population. The MPS VI Modifier is an asymptomatic variant that is contributing to a mild phenotypic expression of disease in compound heterozygotes (54–56), suggesting selecting against the c.1558G>A variant is not justified or recommended, as it would also reduce the genetic variation in the breed.

The appearance of the cat is influenced by various genes which are often monitored by genetic testing to inform breeding pair selection. In this study, all cats were tested for 26 appearance-associated variants. Information was provided on the frequency at which the trait variants are encountered across breeds, to explain the observed phenotypes. The same trait variants influencing coat color/type and morphology are highly frequent in cats of various breed backgrounds, providing evidence of likely causality. However, we note that the *KIT* gene variant associated with breed-characteristic gloves (white feet) phenotype in Birman cats (58), is observed in various individuals of other breeds without gloves phenotype. Moreover, while most of the trait phenotypes were explained by the known variants, the previously discovered variants associated with shortened tail, extra digits and hairlessness could only explain the presence of some of these phenotypes.

Our analysis of genetic diversity in cat breed populations shows a wide range of diversity levels within and between breeds. We found evidence that, as expected, more recently formed breeds with a more significant number of founding individuals and breeds allowing continued outcrossing tend to have the greatest diversity levels. Maintaining diversity in closed populations is challenging, and the use of outcrossing may help maintain and potentially increase diversity levels if widely adopted. The importance of preserving diversity for health and vigor has been widely documented (70–72). Additionally, the disease-associated variant findings in the non-pedigreed cat samples set demonstrate the significance of genetic screening for known disease-associated variants as well.

In conclusion, we demonstrate that several feline disease-associated variants are more widespread across cat breeds and varieties than previously reported, with both dominant and recessive Mendelian disease-associated variants observed in additional breeds and often at higher allele frequency than the breeds in which they were originally discovered. This, in part, demonstrates the effectiveness of proactive genetic testing, which has reduced disease-associated variant frequencies in notably affected breeds. We have also shown that some disease-associated variants are very rare and limited to specific breeds. We report the prevalence at which the three clinically relevant feline blood types occur within breeds and breed types and provide trait variant frequencies across the feline population. We have combined genotype information with phenotypic information to investigate and re-evaluate causality in different breed backgrounds, confirming causal relationships for some variants and weak evidence of penetrance for other variants. In summary, genetic testing can be used to inform breeding decisions aiming to prevent genetic disease, while a concurrent goal should be to maintain genetic diversity in a breed’s population, helping to sustain the breed. As more cats are genotyped, we will learn more about the feline variant heritage in the broader domestic cat population, leading to improved advice to all cat owners. Direct-to-consumer tests help to further raise awareness of various inherited conditions in cats, provide information that owners can share with their veterinarians, and in time, as more genotypic and phenotypic data are collected, will enable the genetics of common complex feline disease to be deciphered, paving the way for personalized precision healthcare with the potential to ultimately improve welfare for all cats.

## Acknowledgements

We extend our warmest thanks to all the cat owners and breeders who enabled the present study by voluntarily submitting photos and clinical documentation through their interest in advancing feline genetics research; we issue special thanks to the participants in the Wisdom Panel-The International Cat Association (TICA) State of the Cat Study for their early support and enthusiasm for this work. We thank Professor Leslie Lyons from the University of Missouri and various anonymous cat owners for providing validation samples. Excellent advice and support were provided by TICA, Finnish Cat Association, Anthony Hutcherson, Adriana Kajon, Suvi Ruotanen, Dr. Casey A. Brookhart-Knox, Dr. Jason Huff and Dr. Susan Puckett; all of whom are deeply thanked.

## Supporting Material

S1 Table. The summary of 11,036 tested pedigreed and non-pedigreed cats.

S2 Table. Tested disease and trait associated variants.

S3 Table. All tested disease and trait genotype data for 11,036 tested cats.

S4 Table. All tested disease and trait variant frequencies for 11,036 tested cats.

S5 Table. Clustered breed information for representing the proportions of the different blood types with > tested individuals.

S6 Table. Genetic diversity for all breeds with >15 individuals tested.

